# *Arabidopsis* transcriptome responses to low water potential using high throughput plate assays

**DOI:** 10.1101/2022.11.25.517922

**Authors:** Stephen Gonzalez, Joseph Swift, Adi Yaaran, Jiaying Xu, Charlotte Miller, Natanella Illouz-Eliaz, Joseph R. Nery, Wolfgang Busch, Yotam Zait, Joseph R. Ecker

## Abstract

Soil-free assays that induce water stress are routinely used to investigate drought responses in the plant *Arabidopsis thaliana*. Due to their ease of use, the research community often relies on polyethylene glycol (PEG), mannitol and salt (NaCl) treatments to reduce the water potential of agar media, and thus induce drought conditions in the laboratory. However, while these types of stress can create phenotypes that resemble those of water deficit experienced by soil-grown plants, it remains unclear how these treatments compare at the transcriptional level. Here, we demonstrate that these different methods of lowering water potential elicit both shared and distinct transcriptional responses in *Arabidopsis* shoot and root tissue. When we compared these transcriptional responses to those found in *Arabidopsis* roots subject to vermiculite drying, we discovered many genes induced by vermiculite drying were repressed by low water potential treatments on agar plates (and vice versa). Additionally, we also tested another method for lowering water potential of agar media. By increasing the nutrient content and tensile strength of agar, we show the ‘hard agar’ (HA) treatment can be leveraged as a high-throughput assay to investigate natural variation in *Arabidopsis* growth responses to low water potential.

## Introduction

As climate change advances, improving crop drought tolerance will be key for ensuring food security (*1, 2*). This has led to intense research at the molecular level to find novel loci and alleles that drive plant responses to drought conditions. Such investigations benefit from simple assays that can reproduce drought phenotypes at both the physiological and molecular levels. While some researchers use soil-based assays, these are cumbersome. For example, extracting intact root systems from the soil is difficult, and reproducing the rate at which water evaporates from the soil can be challenging (*3*). In light of this, chemical agents such as polyethylene glycol (PEG), mannitol, or salt (NaCl) are often employed to induce drought stress. When present in aqueous or agar media, they allow precise and dose-dependent control of water potential (*4, 5*). When exposed to these media types, plants exhibit the hallmarks of drought physiology, such as reduced growth rate, reduced stomatal conductance, and increased leaf senescence (*4, 6, 7*). While each of these methods lower water potential and thus induce a drought stress, each method exerts additional and distinct effects. For example NaCl not only induces osmotic stress, but can cause salt toxicity (*6*). Since it is not metabolized by most plants mannitol is considered less toxic (*3*), however evidence suggests it may act as a signaling molecule (*5, 8*). Since both NaCl and mannitol can enter the pores of plant cell walls (*9, 10*), they can induce plasmolysis, a process that does not typically occur under mild water deficit (*9*). Due to its higher molecular weight, PEG treatment avoids this, and instead causes cytorrhysis (*9*), a physiology more common under drought settings (*10, 11*).

The unique impacts PEG, mannitol and NaCl have on plant physiology may also extend to the level of gene expression. Indeed, a broad spectrum of transcriptional changes are documented in response to low water potential, which may be attributed to the specific method employed (*4, 12, 13*). Here, we examine the transcriptional responses to PEG, mannitol and NaCl in *Arabidopsis*, and compare these responses to those elicited when plants are exposed to vermiculite drying. Furthermore, we explore a new approach for reducing water potential.

### Comparing differential gene expression responses elicited by PEG, mannitol and NaCl treatment to vermiculite drying

To understand the impact PEG, mannitol and NaCl treatment have on gene expression, we first tested their physiological effects across a range of doses. To this end, we grew *Arabidopsis* seedlings on agar plates supplemented with Linsmaier & Skoog (LS) nutrients for 14 days on three different doses of each stress type. Dose ranges were chosen based on published literature, and ranged from mild to severe stress levels (*3, 4, 11*). As the dose of each stress type increased, the media’s water potential significantly decreased in a dose-dependent manner (Pearson, *p* < 8.5 × 10^-5^). Across the doses tested, we found that each stress type’s impact on water potential was not statistically different from one other (ANCOVA post-hoc, *p* > 0.05). In response to these treatments, we found shoot biomass significantly decreased in a dose-dependent manner (Pearson, *p* < 2 × 10^-6^) (**Figure 1A - C, Figure 1-figure supplement 1, Supplementary File 1**).

**Figure 1.**
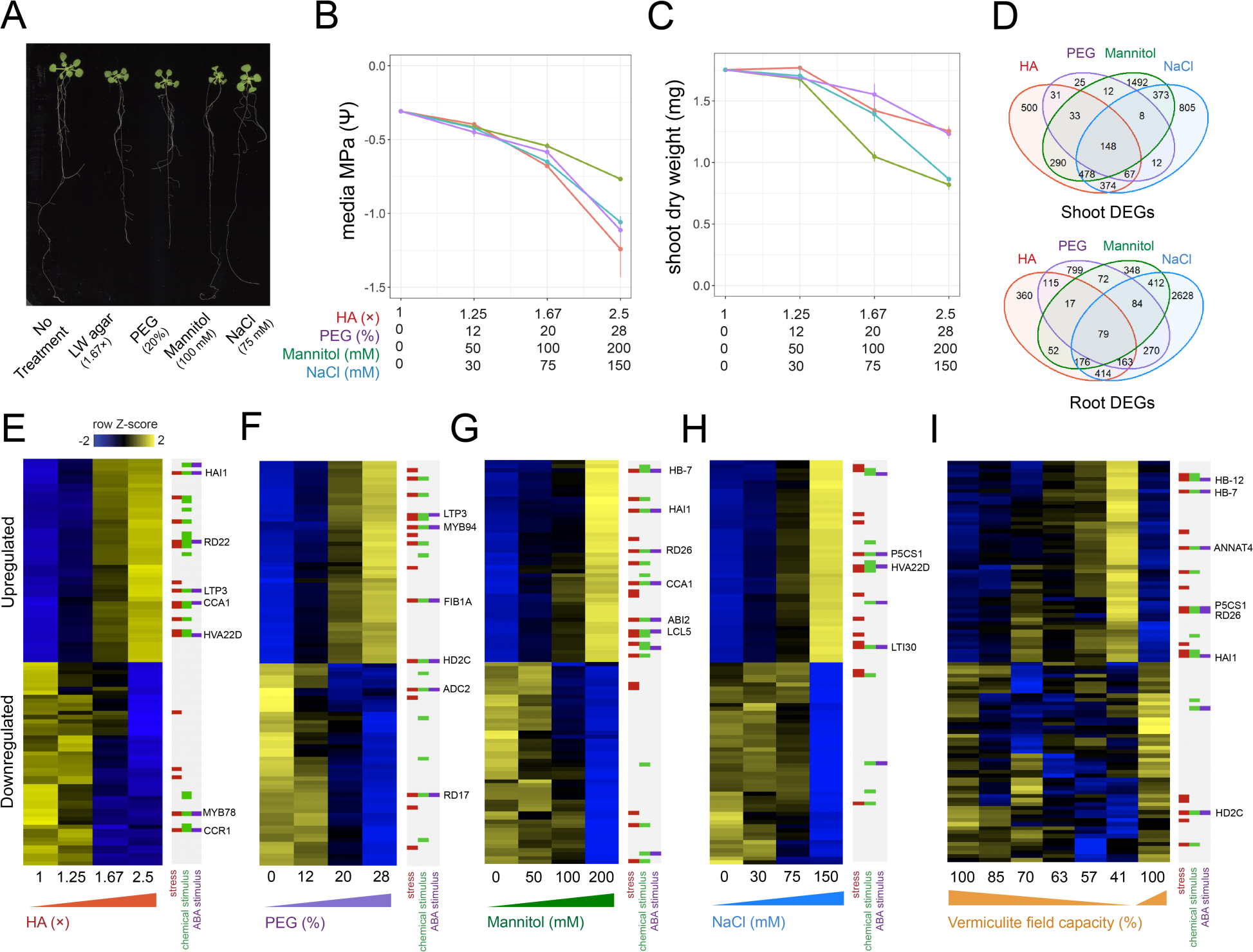
Benchmarking the impact different stress assays have on *Arabidopsis* gene expression. **A:** 22-day-old *Arabidopsis* growth on plates under either 1.67ξ hard agar (HA), 20 % PEG, 100 mM mannitol, or 75 mM NaCl treatments. **B**: Water potential measurements of treatment media (n = 3 - 4). **C:** Dry weight of 22-day-old *Arabidopsis* seedlings under different doses of each stress treatment (n = 11 - 12). **D:** Number and intersect of differentially expressed genes (DEGs) that are dose-responsive to each stress treatment within root and shoot tissue. **E - I:** Heatmaps displaying the top 50 most significant upregulated or downregulated genes in response to **(E)** HA, **(F)** PEG, **(G)** mannitol, **(H)** NaCl and **(I)** vermiculite drying in the *Arabidopsis* root (n = 2 - 3 biological replicates). Membership of Gene Ontology (GO) Terms for ‘response to stress’, ‘response to chemical stimulus’ or ‘response to ABA stimulus’ are indicated.

Genes that change their expression in response to an environmental signal often do so in a dose-responsive manner (*4, 14*). In light of this, we sought to discover genes whose expression was dose-responsive to the amount of PEG, mannitol, or NaCl applied. By identifying stress-responsive genes across a range of doses, we ensured such genes responded to and were directional with the stress as a whole and not induced or repressed at an individual dose. Taking this approach, we sequenced root and shoot bulk transcriptomes by RNA-seq, and associated each gene’s expression with the dose of stress with a linear model. To ensure we captured steady-state differences in gene expression, and avoided trainsent ones, we sequenced root and shoot transcriptome profiles after 14 days of stress exposure (*3, 6, 15*). By these means, we found hundreds of genes that were dose-responsive to each treatment within root and shoot tissue (**Figure 1E - 1H, Figure 1-figure supplement 2, Supplementary File 2**) (adj. *p* < 0.05). We found that a portion of these dose-responsive genes were shared across treatments, suggesting a common response to low water potential (**Figure 1D, Figure 1-supplement Figure 3**). Conversely, we also found a portion of dose-responsive genes were unique to each stress type.

Next, we wanted to compare these different methods of lowering water potential to a pot-based water deficit assay. To perform this experiment, we subjected mature *Arabidopsis* plants grown in pots on vermiculite supplemented with LS media to mild water stress by withholding water for 5 days. During this period, field capacity was reduced from 100 % to 41 %. This treatment led to a reduction in plant biomass (*p* = 1.8 × 10^-3^), as well as seed yield (*p* = 1.2 × 10^-4^), but did not induce visible signs of senescence or wilting (**Figure 1-figure supplement 4, Supplementary File 3**). We assayed root and shoot gene expression responses daily during water loss by RNA-seq. We observed a dose-dependent relationship between a decrease in field capacity and gene expression responses in both roots and shoots, identifying 1,949 differentially expressed genes in roots and 1,792 in shoots (adj. *p* < 0.01) (**Figure 1I, Figure 1-figure supplement 2, Supplementary File 2**). We ensured these genes’ expression patterns recovered upon rewatering (**Figure 1I**). We note that while vermiculite has greater aeration than soil, we found that the genes differentially expressed in roots in response to vermiculite drying largely agreed with a previous report detailing transcriptional responses to soil drying (*16*) (**Figure 2-figure supplement 1**). We also found differentially expressed genes responsive to vermiculite drying agreed with those responsive to transient treatment with Abscisic Acid (ABA), a stress hormone whose levels rise in response to water deficit (*4*) (**Figure 2-figure supplement 2**).

To assess how PEG, mannitol and NaCl treatments compared to the vermiculite drying response described above, we overlapped genes found differentially expressed in each experiment. For shoot tissue, we found genes that were differentially expressed during vermiculite drying overlapped significantly with genes that were differentially expressed by either PEG, mannitol and NaCl treatments (Fisher test, adj. *p* < 0.05). Additionally, there was 88 – 99 % directional agreement within these overlaps, indicating that genes induced or repressed by vermiculite drying were similarly induced or repressed by low water potential treatments on agar (**Figure 2A** & **2B**). Along these lines, across all conditions we saw differential expression of the desiccation-associated genes *RESPONSE TO DESSICATION 20;29B (RD20;29B)* (*17, 18*), the osmo-protectant gene *DELTA1-PYRROLINE-5-CARBOXYLATE SYNTHASE 1* (*P5CS1*), and ABA signaling and biosynthesis genes *HOMEOBOX 7* (HB7) (*19*) and *NINE-CIS-EPOXYCAROTENOID DIOXYGENASE* (*NCED3)* (*20*) (**Figure 2-figure supplement 3**). We note that while we observed agreement in the direction of gene expression across assays, there were differences in the amplitude of gene expression (**Figure 2-figure supplement 3**). This may be due to confounding factors, such as differences in the ranges of water potential tested (**Figure 1B**), or through comparing seedlings grown on plates with mature *Arabidopsis* plants grown on vermiculite.

**Figure 2.**
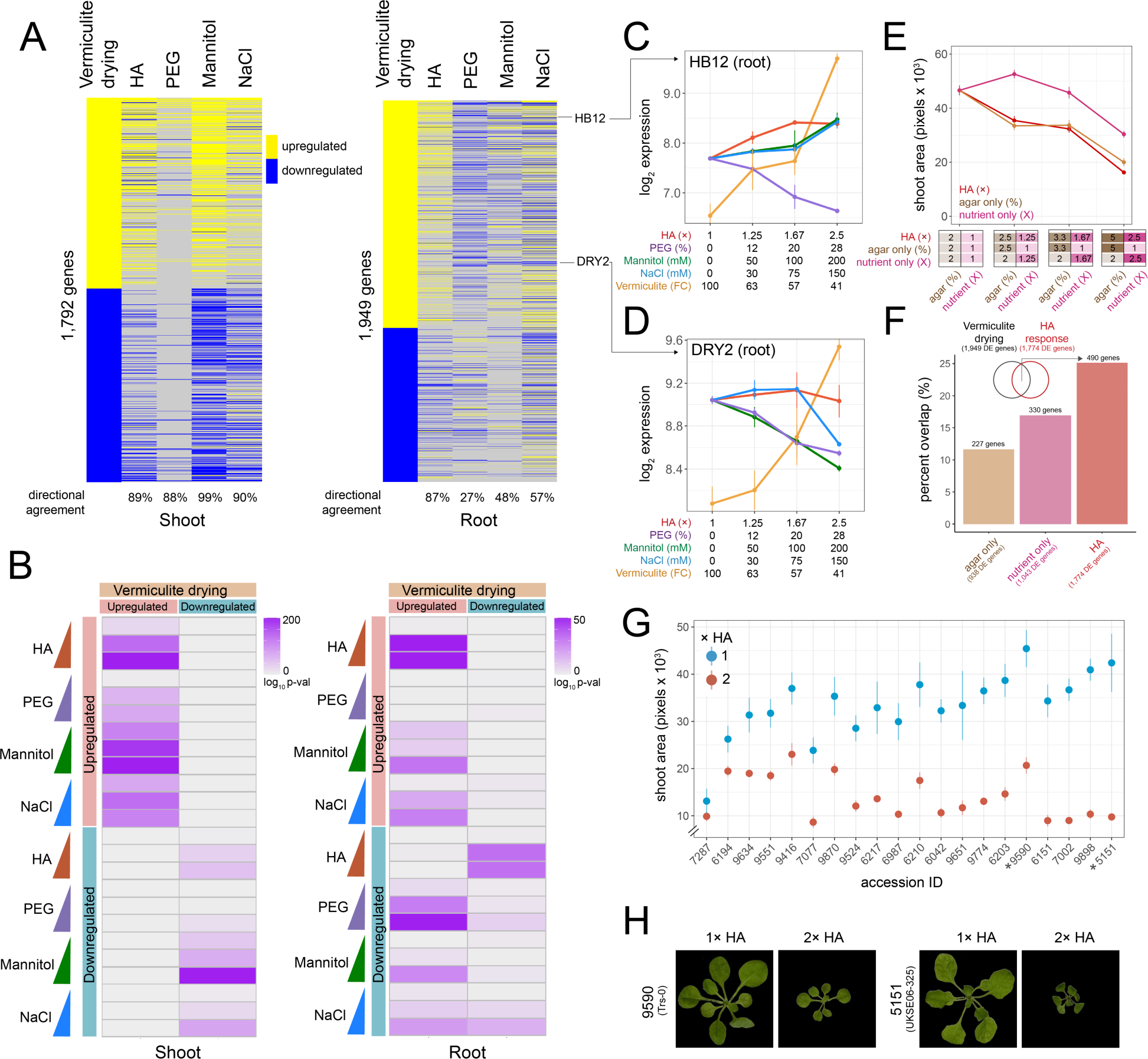
– Comparing HA, PEG, mannitol, and NaCl gene expression responses to vermiculite drying. **A:** Heatmap displaying genes differentially expressed in response to vermiculite drying in shoot or root tissue compared to their dose-responsive expression within each plate-based assay. Level of ‘directional agreement’ (i.e., differentially expressed in the same direction) found within each assay reported. **B:** Overlap analysis of genes found differentially expressed due to vermiculite drying, compared to those found differentially expressed within each dose of PEG, mannitol, NaCl or HA agar assays in either shoot and root. **C - D:** Expression patterns of *HOMEOBOX12* (*HB12)* and *DROUGHT HYPERSENSITIVE 2* (*DRY2)* across each assay in root tissue. **E:** Shoot area of seedlings grown under increasing doses of HA, agar or nutrient concentrations (n = 19). **F:** Number and percent overlap of genes found differentially expressed in response to increasing doses of HA, agar or nutrient concentrations with those differentially expressed in response to vermiculite drying. **G:** Total shoot area of *Arabidopsis* accessions grown under either 1ξ or 2ξ HA agar treatment (n = 5 - 12). **H:** Images of *Arabidopsis Trs-0* or *UKSE06-325* accessions grown on either 1ξ or 2ξ HA treatment.

In root tissue, we found greater variability in transcriptomic responses to the different methods of lowering water potential. In particular, we found a number of genes that were upregulated during vermiculite drying were downregulated by PEG, mannitol and NaCl treatments (and vice versa) (**Figure 2A**). This trend persisted when we assessed genes found differentially expressed at each discrete dose of stress (**Figure 2B**). For example, 27 % of PEG dose-responsive genes shared the same direction of expression seen in vermiculite drying responses. We note that previously published PEG transcriptome datasets largely agreed with our own (**Figure 2-figure supplement 1**). Such differential regulation in comparison to vermiculite drying is exemplified by the expression of genes such as *HOMEOBOX 12* (*HB12*) (*19*) (**Figure 2C**), *GRC2-LIKE 1* (*GCL1*) (*21*), and *RESPONSE TO DEHYDRATION* 21 (*RD21*) (*22*) (**Figure 2-figure supplement 3**). We found genes downregulated by PEG are over-represented in the ‘monooxygenase activity’, and ‘oxygen binding’ Gene Ontology (GO) Terms (*p* < 1 × 10^-15^, **Supplementary File 4**). Mannitol and NaCl held a 48 % and 57 % agreement in gene expression direction with vermiculite drying respectively. Examples of genes that followed this pattern of differential regulation in mannitol and NaCl treatments were *DROUGHT HYPERSENSITIVE 2* (*DRY2*) (*23*) (**Figure 2D**) and *ROOT HAIR SPECIFIC 18* (RHS18) (*24*). NaCl-responsive GO Terms included a specific downregulation of ‘phosphorous metabolic processes’ (*p* = 5.2 × 10^-6^), suggesting that the roots were changing phosphate levels in response to NaCl, a process known to help maintain ion homeostasis (*25*). For mannitol, we observed a specific downregulation of ‘cell wall organization or biogenesis’ and ‘microtubule-based processes’ (*p* < 7.8 × 10^-3^) (**Supplementary File 4**).

### Examining differential gene expression responses to ‘hard agar’ (HA) treatment

In addition to examining PEG, mannitol and NaCl transcriptional responses, we also tested a new way of lowering water potential on an agar plate. We hypothesized that we could induce stress by increasing both the agar and nutrient concentration. We called this media ‘hard agar’ (HA). By testing three different doses (1.25×, 1.67× and 2.5× fold increase in both agar and LS concentration, where 1.0× is 2 % agar and 1 X LS), we found that it limited plant shoot dry weight and media water potential in a similar way to PEG, mannitol and NaCl treatment (**Figure 1A - C**). Additionally, we found that HA treatment limited *Arabidopsis* primary root growth rate, shoot water potential, and photosynthesis efficiency (**Figure 2-figure supplement 4**). At the molecular level, RNA-seq revealed 1,376 and 1,921 genes that were dose-responsive to the level of HA stress in roots and shoots respectively (**Figure 1E & Figure 1-figure supplement 2**). We found that these gene expression responses overlapped significantly with those found differentially expressed in response to vermiculite drying (Fisher test, *p* < 1 × 10^-32^, 87 % directional agreement) (**Figure 2A** & **2B**). HA’s impact can be seen in the gene expression responses of *HB12* (**Figure 2C**), *GCL1* and *RD21* (**Figure 2-figure supplement 3**).

An increase in nutrient concentration can induce salt-like stress while increasing agar concentration will increase tensile stress (*26*). We tested each of these variables separately to understand the role each played in eliciting the gene expression responses found in the HA assay. To do this, we repeated our HA dose experiment, but now increasing only the concentration of LS nutrients (1, 1.25, 1.67 and 2.5 X) or the concentration of agar (2, 2.5, 3.3 and 5 %) (**Figure 2E**). We found a significant decrease in shoot area size in response to an increase in nutrient concentration (Pearson *p* = 1.7 × 10^-7^) and agar concentration (*p* = 3.9 × 10^-13^), where the latter more closely phenocopied the effect of HA (**Figure 2E, Figure 2-figure supplement 5, Supplementary File 5**). Since the increase in nutrient concentration alone was responsible for changing media water potential, the phenotypic response to increased agar concentration was not in response to a lower water potential (**Figure 2-figure supplement 5**). Next, we examined the transcriptional responses underlying nutrient and agar responses by sequencing root tissue across each dose tested. Through linear modeling, we found 1,043 genes and 938 genes that were dose-responsive to nutrient or agar concentration, respectively. Then, we investigated how these genes compared to those found in vermiculite drying responses. We found that genes differentially expressed in response to an increase in agar or nutrient concentration overlapped 12 % and 17 % of vermiculite drying responsive gene expression respectively (permutation test, *p* < 0.05) (**Figure 2F, Figure 2-figure supplement 5, Supplementary File 6**). However, we found genes differentially expressed in response to HA treatment led to a higher overlap (26 %), suggesting that both nutrient and agar concentration contribute to the similarity between HA treatment and vermiculite drying.

Finally, we tested if our HA assay was sensitive enough to detect phenotypic variability. To achieve this, we grew 20 different *Arabidopsis* ecotypes on 2ξ HA agar (4% agar, 2 X LS), where ecotypes were selected from a previous drought study that assessed fitness in a common garden experiment (*27*). By comparing the total shoot area after three weeks of growth, we found that our assay revealed variability in shoot growth responses (**Figure 2G** & **2H, Supplementary File 7**). Furthermore, we found that the greater the impact HA agar had on reducing an accession’s relative shoot size, the better the accession’s fitness was, as measured under field conditions (*27*) (Spearman *p* = 0.04, **Figure 2-figure supplement 6**). This is likely because smaller shoot systems have a better chance of survival and reproduction (*28*). This suggests that our assay may be useful for screening for novel drought-associated loci among a wider group of accessions or mutants (*29, 30*).

In summary, our investigation has assessed the shared and unique impacts of agar-based low water potential treatments on gene expression. We also compared these effects with the expression patterns elicited during vermiculite drying. We found each plate-based assay generated similar responses in shoot tissue, but more varied responses in root tissue. We note that our comparative analysis focuses largely on transcriptomic responses in *Arabidopsis.* We suggest investigating gene expression responses in other species as future work. Here, we also introduce another method for lowering water potential. By increasing nutrient and agar concentration, our HA approach also induced gene expression responses comparable to vermiculite drying. We describe how to make this media within the **Materials and Methods**.

## Supplementary

**Supplementary File 1** – Plant physiological measurements

**Supplementary File 2** – Differentially expressed genes and normalized counts in HA, PEG, Mannitol, NaCl or vermiculite drying experiments

**Supplementary File 3** – Vermiculite drying assay measurements

**Supplementary File 4** – GO Term enrichment of differentially expressed genes

**Supplementary File 5** – Shoot area of seedlings grown under different agar and nutrient concentrations

**Supplementary File 6** – Differentially expressed genes and normalized counts in response to changes in nutrient or agar concentration

**Supplementary File 7** – Shoot area of different *Arabidopsis* accessions grown on HA media

## Acknowledgements

We thank Renee Garza for critical reading of the manuscript. J.S is an Open Philanthropy awardee of Life Science Research Foundation, as well as recipient of the Pratt Industries American-Australian Association Scholarship. J.R.E is an Investigator of the Howard Hughes Medical Institute.

## Data Availability

Raw sequencing data is available at the National Center for Biotechnology Information Sequence Read Archive (accession number PRJNA904764). Normalized read counts and raw phenotypic datasets can be found in the Supplementary Material.

## Materials and Methods

### Hard Agar Assay

*Arabidopsis* seedlings were grown on vertical plates for 8 days under short day conditions (8 h light, 21 C, 150 umoles light) on agar media (1 × Linsmaier & Skoog (LS) (Cassion LSP03) media, 1% sucrose, 2% agar, pH 5.7). We note LS media is identical to Murashige & Skoog (MS) media in inorganic salt content, but lacks Glycine, Nicotinic Acid and Pyridoxine HCl. After 8 days, plants were transferred to ‘hard agar’ (HA) plates. The 1.0× plate consisted of 2 % and 1 X LS media, with no sucrose (pH 5.7) at a final volume of 75 mL. Subsequent doses of increased nutrient and agar concentration (1.25×, 1.67× and 2.5× fold increase) were made by preparing the same media but reducing the amount of water present. For example, the 1.25×, treatment plate contained 60 mL of 2.5% agar and 1.25× LS media. We note that the volume of HA itself has minimal impact gene expression responses (**Figure 2-figure supplement 7**). On day 14, 2 hours after subjective dawn, shoot and root samples were flash frozen (6 plants per replicate). In total, we collected 16 samples for RNA-seq analysis (2 organs, 2-3 biological replicates, 3 treatment levels). We also collected a non-treated control set (2 biological replicates).

To test different *Arabidopsis* accessions on HA agar, plants were sown on either 1×, or 2× treatments as described above, however supplemented with 0.5% or 1% sucrose respectively to encourage germination. Seedlings were grown for 3 weeks under short day conditions in before imaging plates in duplicate (n = 2 - 5 plants per plate) (**Supplementary File 7**). Shoot area was calculated from images using Plant Growth Tracker (GitHub - https://github.com/jiayinghsu/plant_growth_tracker).

### Vermiculite Drying Assay

*Arabidopsis* seedlings were grown on vertical plates for 17 days under short day conditions (8 h light, 21 C, 150 umoles light) on agar media (1×LS, 1% sucrose, 2% agar, pH 5.7), before transfer to vermiculite saturated with 0.75 × LS media. We note at the timing of transfer lateral root formation had begun. Plants were then grown on vermiculite at 100% field capacity (FC) for 12 days (8 h light, 21 C, 150 umoles light). On the 13^th^ day, the first time point was sampled (4.5 hours after subjective dawn) where tissue was flash frozen in liquid nitrogen. After this, excess aqueous solution was drained from each pot, and then each pot was calibrated to 1 × FC. Plant tissue was harvested each day on subsequent days at the same time of day. Each day, pots were weighed to measure extent of evaporation. By these means, FC was measured (**Figure 1-figure supplement 4**). After the 5^th^ day sample was taken, water was re-added to the remaining pots to an excess of 1× FC. ∼ 15 plants were sampled per time point. In total, we harvested 78 tissue samples for RNA-seq (3 biological replicates, 2 organ types, 7 days, 2 treatments). Plants were then left to grow under long day conditions until flowering. Seeds were harvested, dried, and weighed (n = 50 plants per treatment).

### Polyethylene Glycol (PEG) Stress Assay

*Arabidopsis* seedlings were grown on vertical plates for 8 days under short-day conditions (8 h light, 21 C, 150 umoles light) on agar media (1×LS, 1% sucrose, 2% agar, pH 5.7), before transfer to polyethylene glycol (PEG) media of varying concentrations. PEG media plates were prepared by dissolving crystalline 6000 MW PEG into freshly autoclaved 1× LS media pH 5.7 and pouring 50 mL of PEG media solution onto 1× LS, 2% agar, media plates (pH 5.7), letting the PEG solution diffuse into the solid media overnight, then pouring off excess and transferring seedlings to PEG infused media plates as described in (*11*). Plants were grown under 3 different treatments (12%, 20%, and 28% PEG solution w/v) for 14 days. On day 14, 2 hours after subjective dawn, shoot and root samples were flash frozen (6 plants per replicate). In total, we collected 16 samples for RNA-seq analysis (2 organs, 2-3 biological replicates, 3 treatment levels).

### Mannitol and NaCl Osmotic Stress Assays

*Arabidopsis* seedlings were grown on vertical plates for 8 days under short-day conditions (8 h light, 21 C, 150 umoles light) on agar media (1× LS, 1% sucrose, 2% agar, pH 5.7), before transfer to either mannitol or salt (NaCl) media of varying concentrations. Mannitol and NaCl media plates were prepared by adding respective volume of stock solution to 1× LS, 2% agar, pH 5.7 media before autoclaving. Plants were grown under 3 different treatments of mannitol or NaCl (50 mM, 100 mM and 200 mM for mannitol, 30 mM, 75 mM, and 150 mM for NaCl) for 14 days. On day 14, 2 hours after subjective dawn, shoot and root samples were flash frozen (6 plants per replicate). In total, for either Mannitol or NaCl treatment experiments, we collected 18 samples for RNA-seq analysis (2 organs, 3 biological replicates, 3 treatment levels).

### ABA Exogenous Treatment Assay

*Arabidopsis* seedlings were grown on vertical plates for 8 days under short-day conditions (8 h light, 21 C, 150 umoles light) on agar media (1× LS, 1% sucrose, 2% agar, pH 5.7), before transfer to 1× LS, 2% agar, pH 5.7 control media and grown for 14 days. On day 14, abscisic acid (ABA) solutions of 1 uM, 5 uM and 10 uM were prepared from 10 mM ABA dissolved in ethanol stock, as well as a mock treatment solution containing 0.1% ethanol concentration. 30 min after subjective dawn, 15 mL of each solution was dispersed onto the roots of the seedlings. After 1 min of treatment, the ABA solution was removed from the plates, and the plates returned to the growth chamber. 2 hours after subjective dawn, shoot and root samples were flash frozen (6 plants per replicate). In total, we collected 8 samples for RNA-seq analysis (root tissue only, 2 biological replicates, 4 conditions).

### Osmotic Potential Measurements

The water potential of media was determined considering it equivalent to the osmotic potential (Ψs). Osmotic potential was measured using a vapor pressure osmometer (Model 5600, ELITech Group; Puteaux, France). Readings were taken from melted agar media constituted with the particular stress type. Osmolality readings for each sample obtained were converted to megapascals (MPa) using the equation Ψs = -CRT, where C is the molar concentration, R is the universal gas constant, T is the temperature in Kelvin. We note that to measure the water potential of PEG treatment media, we infiltrated the PEG solution into plates as described above, and then melted the PEG-infiltrated agar for measurement with the osmometer. We assessed the osmotic potential of shoot tissue two weeks after transplanting the seedlings to HA agar media, After immersion in liquid nitrogen 3 shoots were placed into 0.5-ml tubes and centrifuged to extract the tissue sap. The osmotic potential (Ψs) of the extracted sap was determined using a vapor pressure osmometer.

### Chlorophyll fluorescence measurements

Chlorophyll fluorescence was assessed in eight seedlings of each plate using the Walz PAM IMAGING PAM M-series IMAG-K7 (MAXI) fluorometer. For every experiment, leaves were pre-conditioned in the dark for 1 h. The maximum quantum yield of PSII (Fv/Fm) was calculated using the formula:

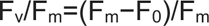

Where Fv is the variable fluorescence, Fm is the maximal fluorescence following 1 h of dark adaptation and F0 is the minimal fluorescence level of a dark-adapted leaf when all photosystem II (PSII) reaction centers are open.

### Root Growth Rate Measurements

*Arabidopsis* seedlings were grown on vertical plates for 8 days under short day conditions (8 h light, 21 C, 150 umoles light) on agar media (1× LS, 1% sucrose, 2% agar), before transfer to 2.5× HA, 28% PEG, 150 mM mannitol or 150 mM NaCl treatment plates as described above. Root images were acquired every two days for a total of 8 days using scanners. Primary root length, defined as the length (scaled to cm) from hypocotyl base to root tip, was quantified using image J. For each treatment we screened 4 plates, with each plate holding 4 individual plants.

### RNA-extraction and Library Preparation

Plant tissue was crushed using the TissueLyser (Agilent) and RNA extracted using RNeasy Mini Kit (Qiagen). Number of biological replicates per library ranged between RNA quality was assessed using Tape station High Sensitivity RNA assay (Agilent). 0.5 - 1 ug of total RNA proceeded to library preparation, where libraries were prepared using TruSeq stranded mRNA kit (Illumina). Resulting libraries were sequenced on the NovaSeq 6000 (Illumina) with 2×150 bp paired-end read chemistry. Read sequences were aligned to the *Arabidopsis* TAIR10 genome using HISAT2 (*31*), and gene counts called using HT-seq (*32*), by relying on Araport11 annotation (*33*). Normalized counts can be found in **Supplementary File 2**. For each organ, libraries from all experiments were normalized together before calling differential expression.

### Statistical Analysis

To detect differential expression in our drought assay on vermiculite, we called differential expression using a linear model using the DESeq2 LRT function to associate a change in field capacity with change in gene expression. The same statistical approach was used to associate a change in a gene’s expression to changes in dose of HA, PEG, mannitol, and NaCl, as well as changes in agar concentration, nutrient concentration and volume of agar used. Resulting model p-values were adjusted to account for false discovery (*p*-value < 0.05). The complete list of differentially expressed genes for each experiment can be found in **Supplementary File 2** and **Supplementary File 6**. Pairwise differential gene expression was called using DESeq2 (*34*). Specifically, for plate based assays, we called differential expression by comparing the control treatment to each treatment dose, using an adjusted *p*-value threshold of 0.05. Overlap analyses were performed using Fisher exact tests, with an adjusted *p*-value threshold of 0.05. The background for these intersects was all expressed genes within the respective organ. Permutation tests and GO Term enrichment analyses was performed in VirtualPlant (*35*), with all expressed genes within the respective organ used as background.

---

**Figure 1 – figure supplement 1.**
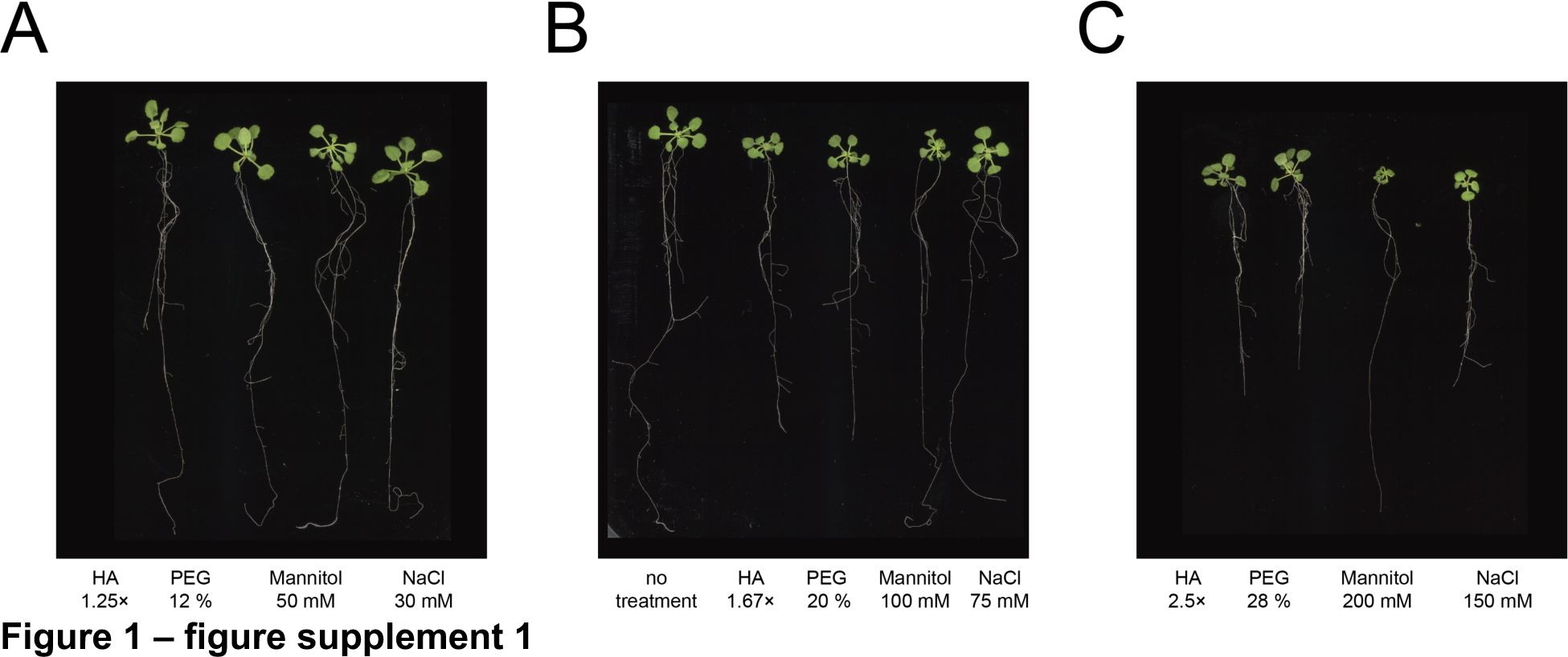
Plant growth responses to stress assays. **A – C:** Images of 22-day-old *Arabidopsis* seedlings grown under different doses of each agar stress assay.

**Figure 1 - figure supplement 2.**
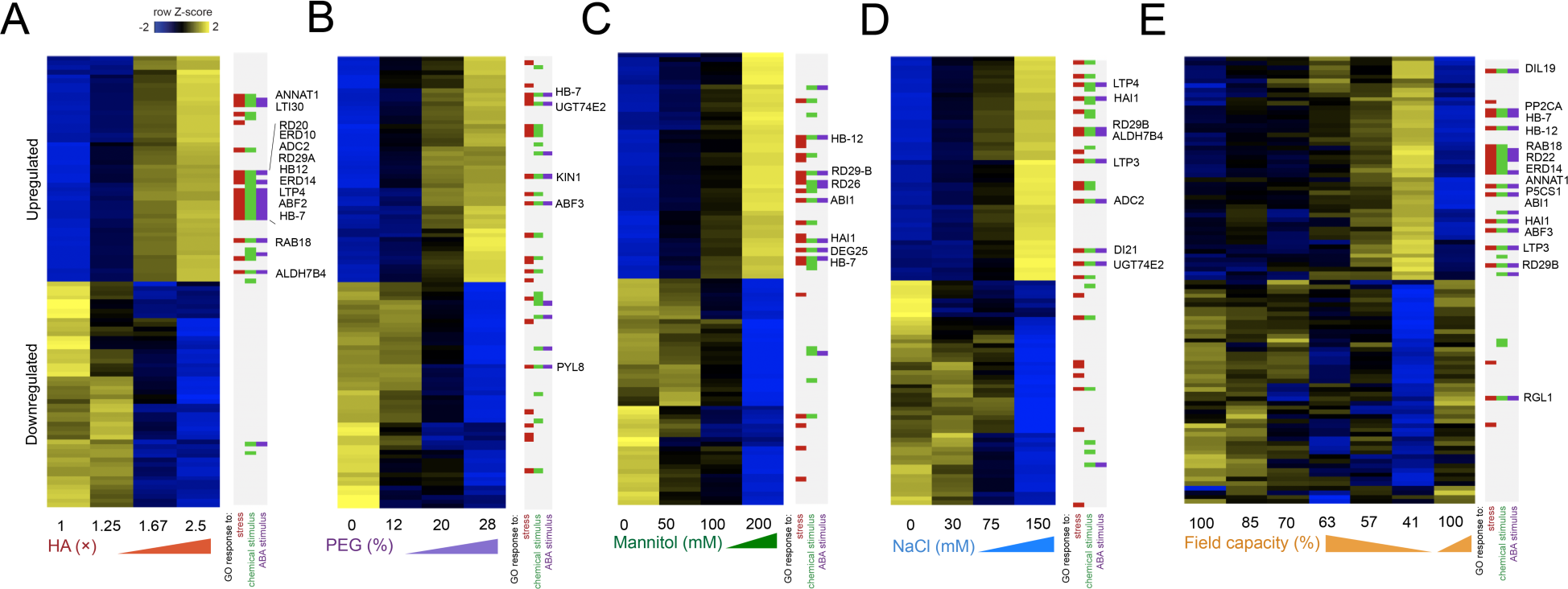
Shoot gene expression responses to each stress assay are dose responsive. Heatmap displaying the top 50 most significant upregulated or downregulated genes in shoots in response to **(A)** HA, **(B)** PEG, **(C)** mannitol, **(D)** NaCl and **(E)** vermiculite drying (n = 2 - 3 biological replicates). Membership of Gene Ontology (GO) Terms for ‘response to stress’, ‘response to chemical stimulus’ or ‘response to ABA stimulus’ indicated.

**Figure 1 - figure supplement 3.**
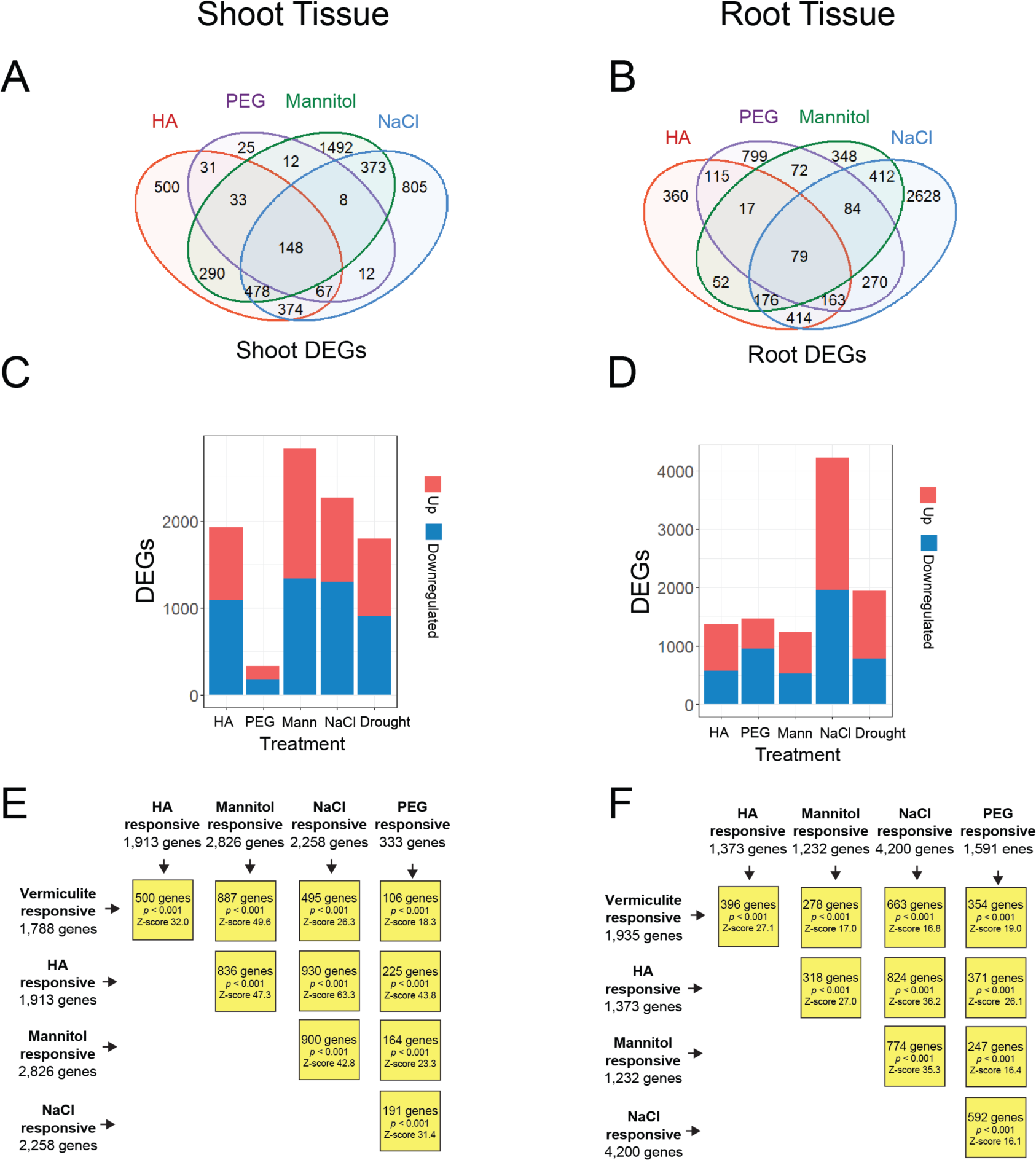
Overlapping differentially expressed genes (DEGs) responsive to different assay types. Overlap of dose-responsive differentially expressed genes in shoot **(A)** and root **(B)** in response to either HA, PEG, mannitol or NaCl (replicated from Figure 1). Number of upregulated or downregulated dose-responsive genes in response to each treatment type in shoot **(C)** and root **(D)**. Overlapping gene sets in **(E)** shoot or **(F)** root tissue (permutation test, p < 0.001).

**Figure 1 - figure supplement 4.**
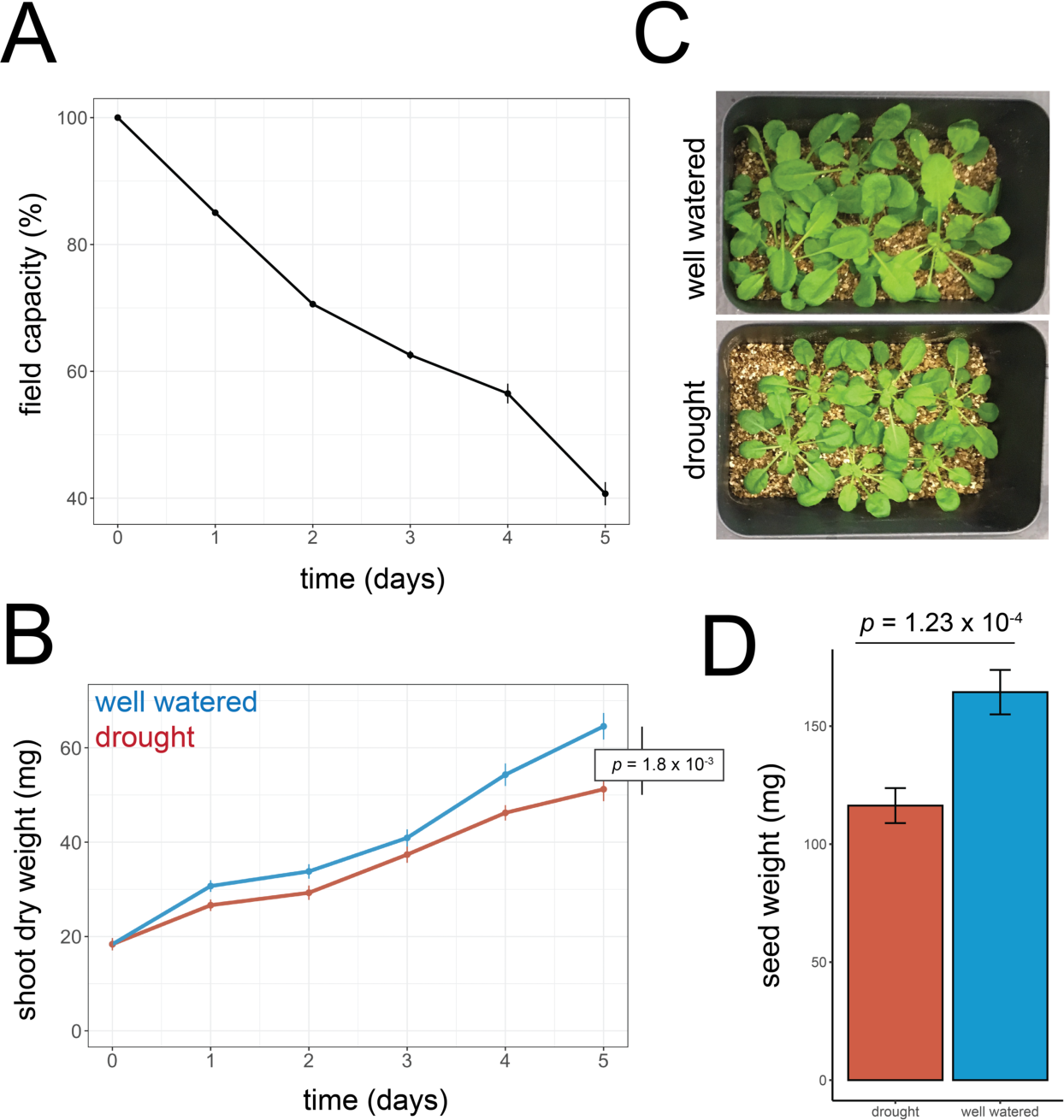
Treating vermiculite-grown *Arabidopsis* plants to mild drought stress. **A:** Field capacity measurements of vermiculite as water evaporated over a 5-day period (n = 6 - 12). **B:** Shoot dry weight of *Arabidopsis* rosettes as they grew either under well-watered conditions or drought conditions over a 5-day period (t-test, n = 12). **C:** Images of plants after 5 days of water stress. **D:** Seed yield resulting from *Arabidopsis* plants after drought recovery (t-test, n = 50).

**Figure 2-figure supplement 1.**
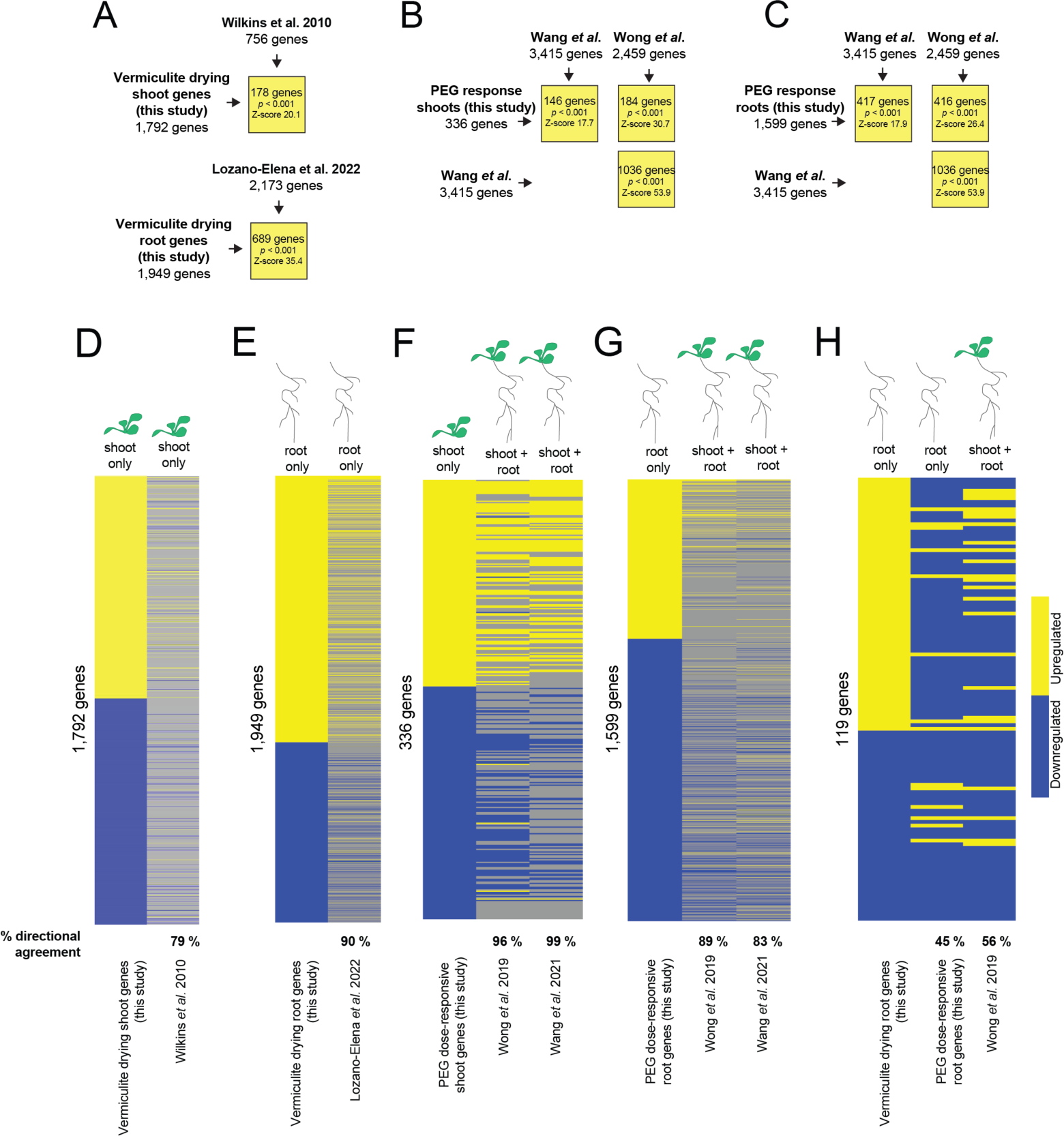
Comparing gene expression response to vermiculite drying and PEG treatment with previous studies. **A:** Intersect analysis of root or shoot genes found differentially expressed in response to vermiculite drying within this study, and genes found differentially expressed in response to soil drying by Lozano-Elena *et al.* (*16*) or by Wilkins *et al.* (*36*) (permutation test, p < 0.001). **B - C:** Intersect analysis of genes found differentially expressed in response to PEG treatment in shoot **(B)** or root **(C)** in this study, with those found differentially expressed in response to PEG treatment by Wong *et al.* (*37*) and Wang *et al.* (*38*) (permutation test, p < 0.001). **D - E:** Heatmap displaying direction of shoot **(D)** or root **(E)** differentially expressed in response to vermiculite drying (this study) and Wilkins *et al.* (*36*) or Lozano-Elena *et al.* (*16*) respectively. Directional agreement with this study’s vermiculite drying response indicated. **F - G :** Heatmap displaying direction of genes differentially expressed in response to PEG treatment across each study. We note that both Wong *et al.* and Wang *et al.* assess transcriptomic responses of whole seedlings (both root and shoot), and thus we compare our shoot **(F)** and root **(G)** data separately. **E:** Examining the 119 genes that were differentially expressed in response to drought (this study), PEG treatment (this study) and PEG treatment reported in Wong *et al*.

**Figure 2-figure supplement 2.**
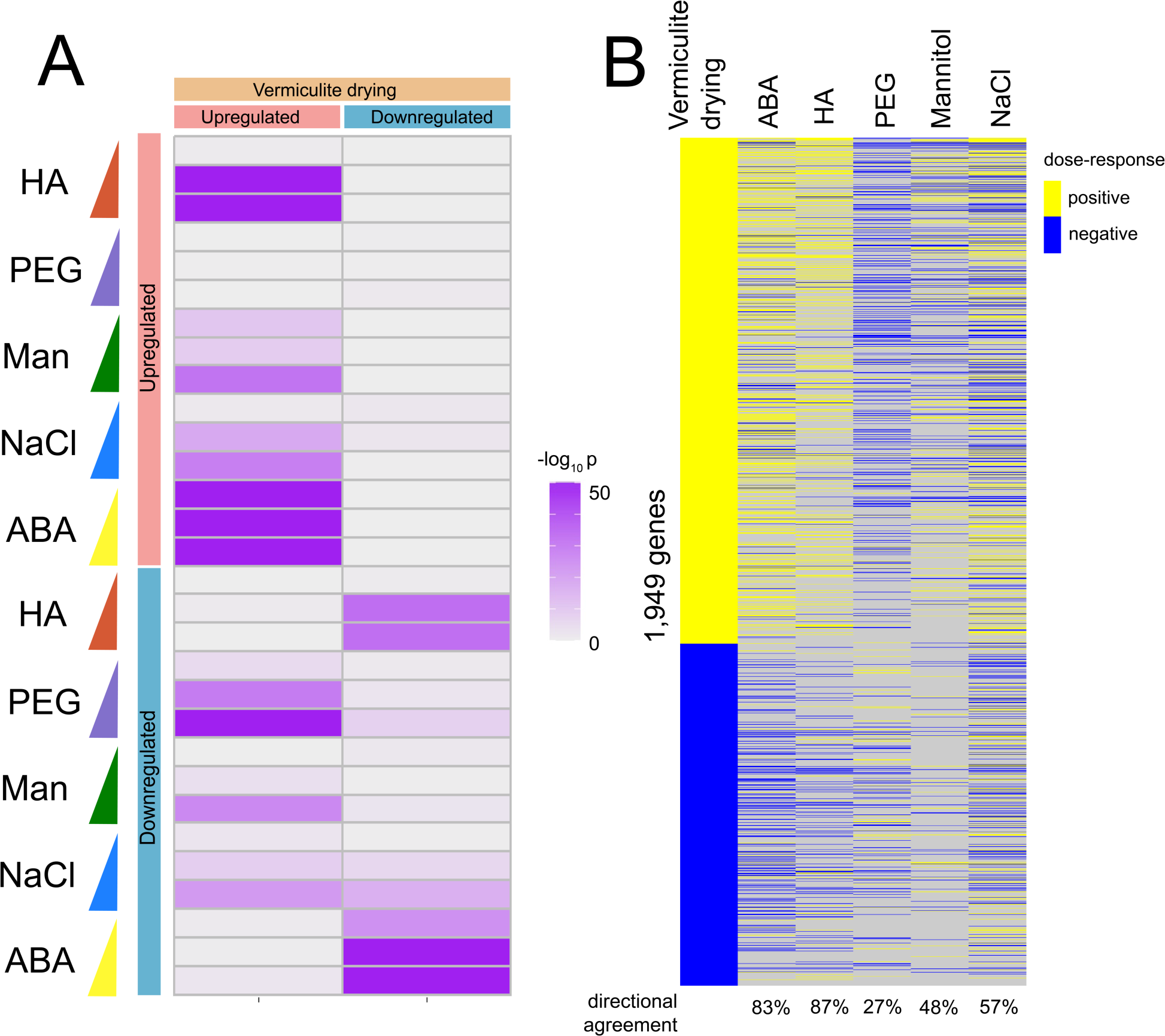
Comparing ABA-induced differential expression to vermiculite drying and HA induced gene expression patterns. **A:** Overlap analysis of genes found differentially expressed in response to vermiculite drying, compared to those within each dose of either transient ABA treatment, PEG, mannitol, NaCl, or HA agar assays in both root and shoot (Fisher exact test, adj. *p* < 0.05). **B:** Heatmap displaying genes differentially expressed under vermiculite drying in root tissue compared to their dose-responsive expression within each stress assay. Direction of gene expression agreement with vermiculite-drying responsive gene expression (i.e., ‘directional agreement’) indicated.

**Figure 2-figure supplement 3.**
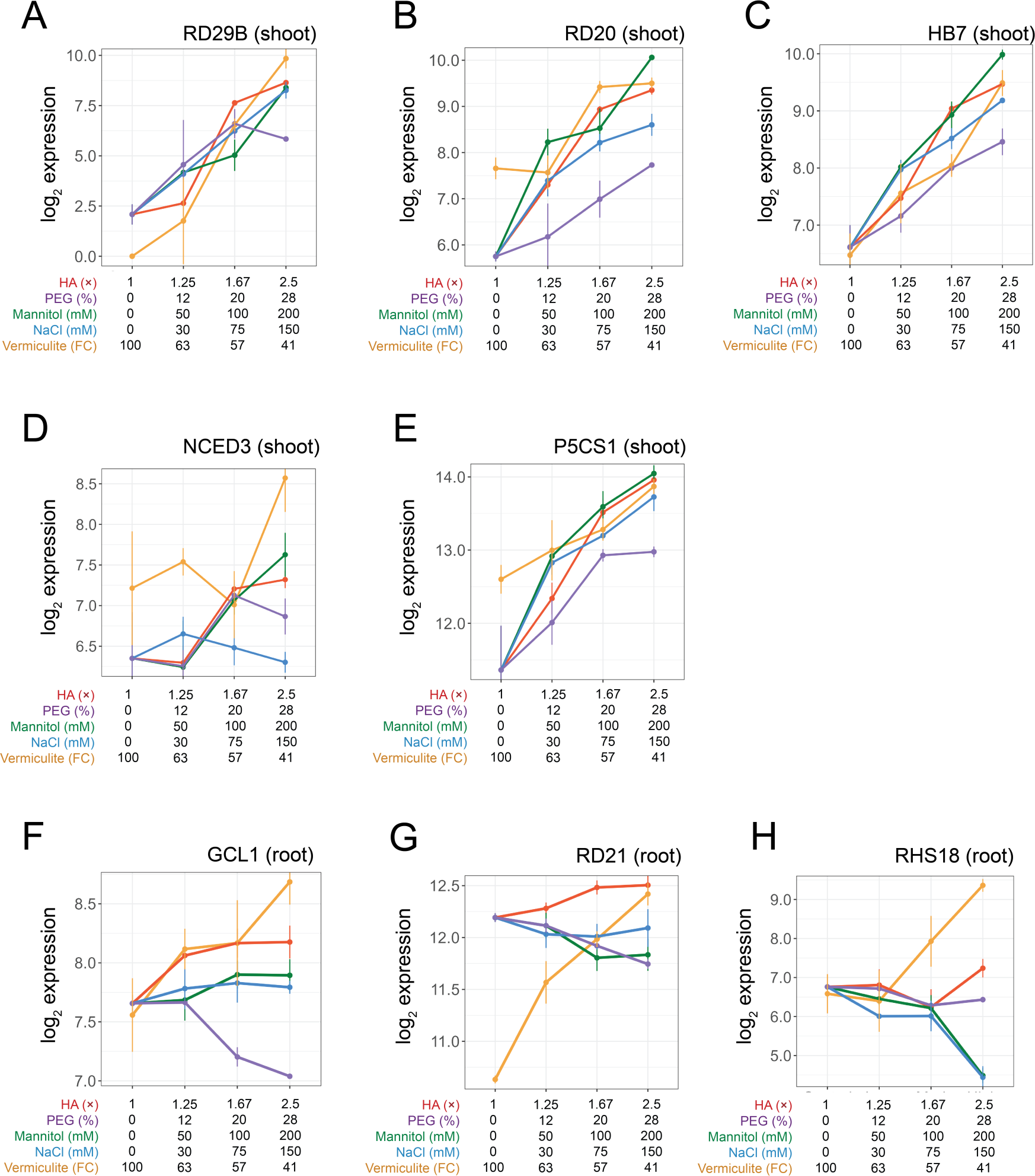
Gene expression profiles of individual genes. **A - E:** Expression patterns of individual genes under doses of each assay in shoot tissue: **(A)** RD29B, **(B)** RD20, **(C)** HB7, **(D)** NCED3 and **(E)** P5CS1. **F - H:** Expression patterns of individual genes under doses of each assay in root tissue: **(F)** GCL1, **(G)** RD21 and **(H)** RHS18.

**Figure 2-figure supplement 4.**
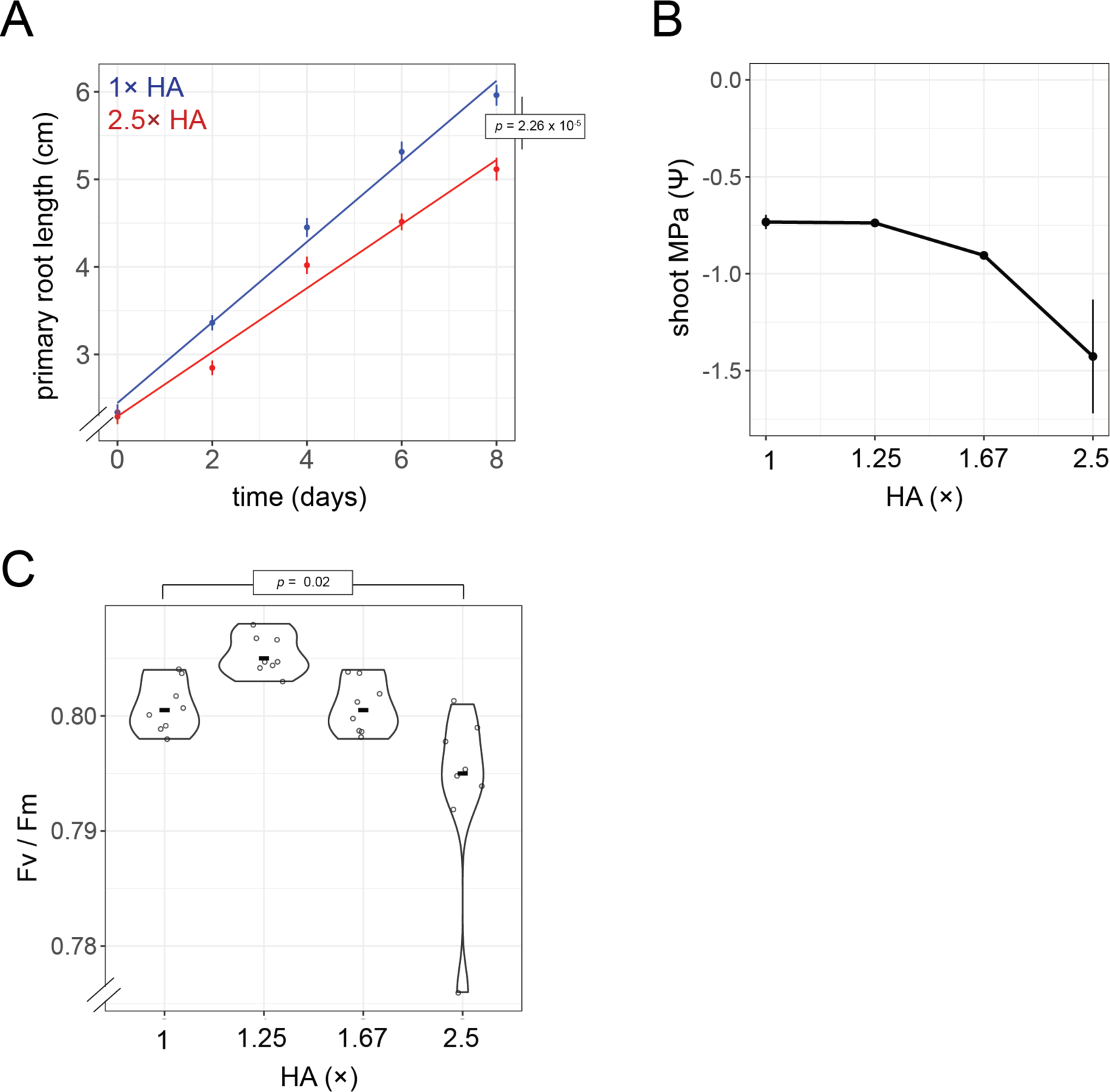
Physiological measurements of *Arabidopsis* seedlings in response to HA treatment. **A:** Measurement of primary root growth rate across 8 days of growth under 1×HA (no treatment) and 2.5× HA conditions (n= 16, t-test *p*). **B**: Shoot water potential measurements of seedlings grown under different HA media doses (n = 3, Pearson *p* = 0.009). **C**: Measurement of maximum quantum yield of PSII (Fv / Fm) under different HA media doses (n=4, t-test *p*).

**Figure 2-figure supplement 5.**
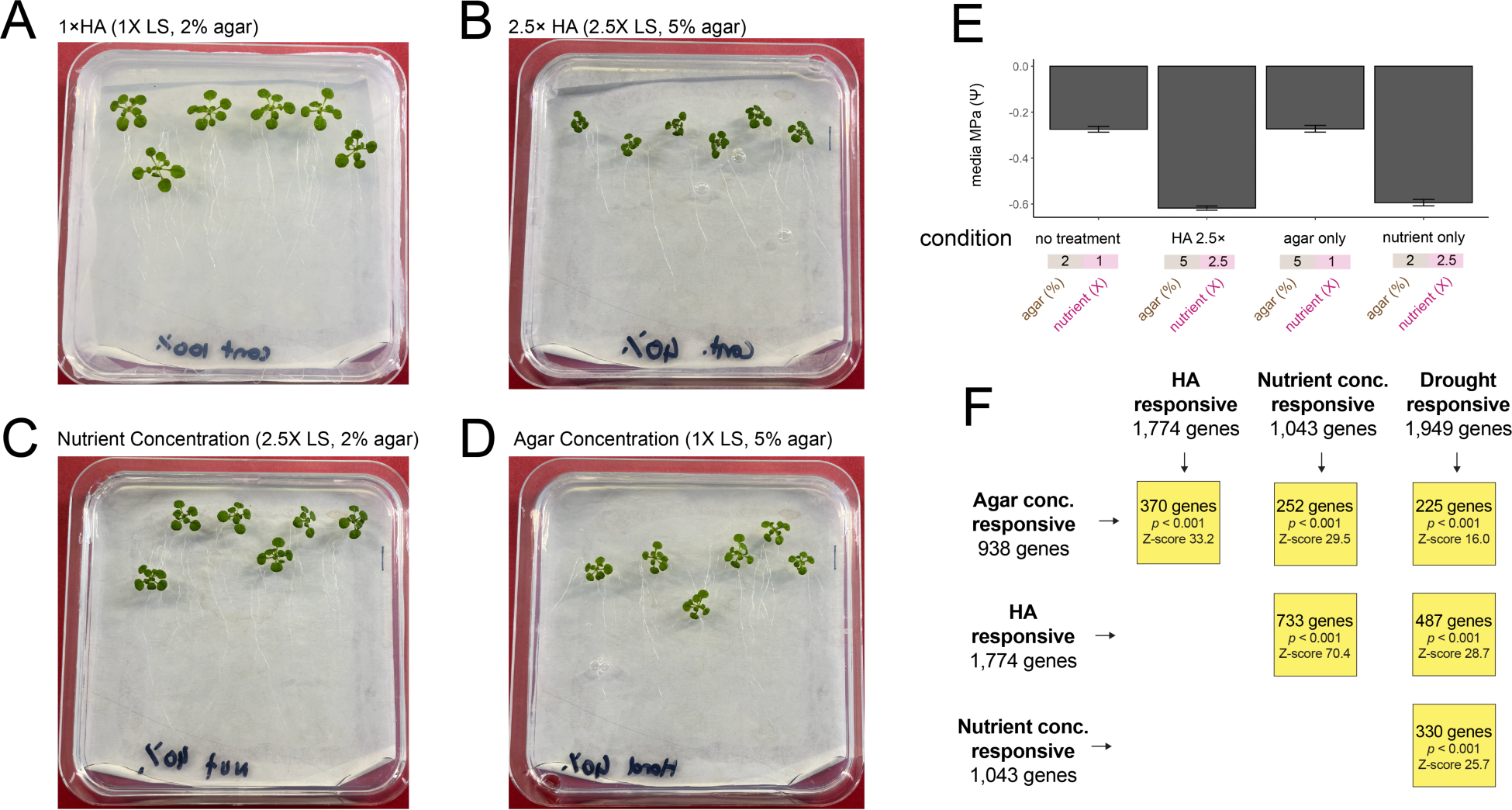
Comparing the separate effects of nutrient concentration and agar concentration on seedling growth. Image of *Arabidopsis* seedlings grown on either **(A)** 1× HA agar (i.e. 1X LS, 2% agar) or **(B)** 2.5ξ HA, which increased both nutrient and agar concentrations to 2.5X and 5%, respectively. **C:** Image of seedlings grown on an increased 2.5X nutrient concentration (without a change in agar concentration). **D:** Image of seedlings grown on an increased 5% agar concentration (without a change in nutrient concentration). **E:** Water potential measurements of media presented in **A – D** (n = 3). **F:** Intersection of differentially expressed genes responsive to either agar concentration, nutrient concentration, HA treatment or drought stress (permutation test, *p* < 0.001).

**Figure 2-figure supplement 6.**
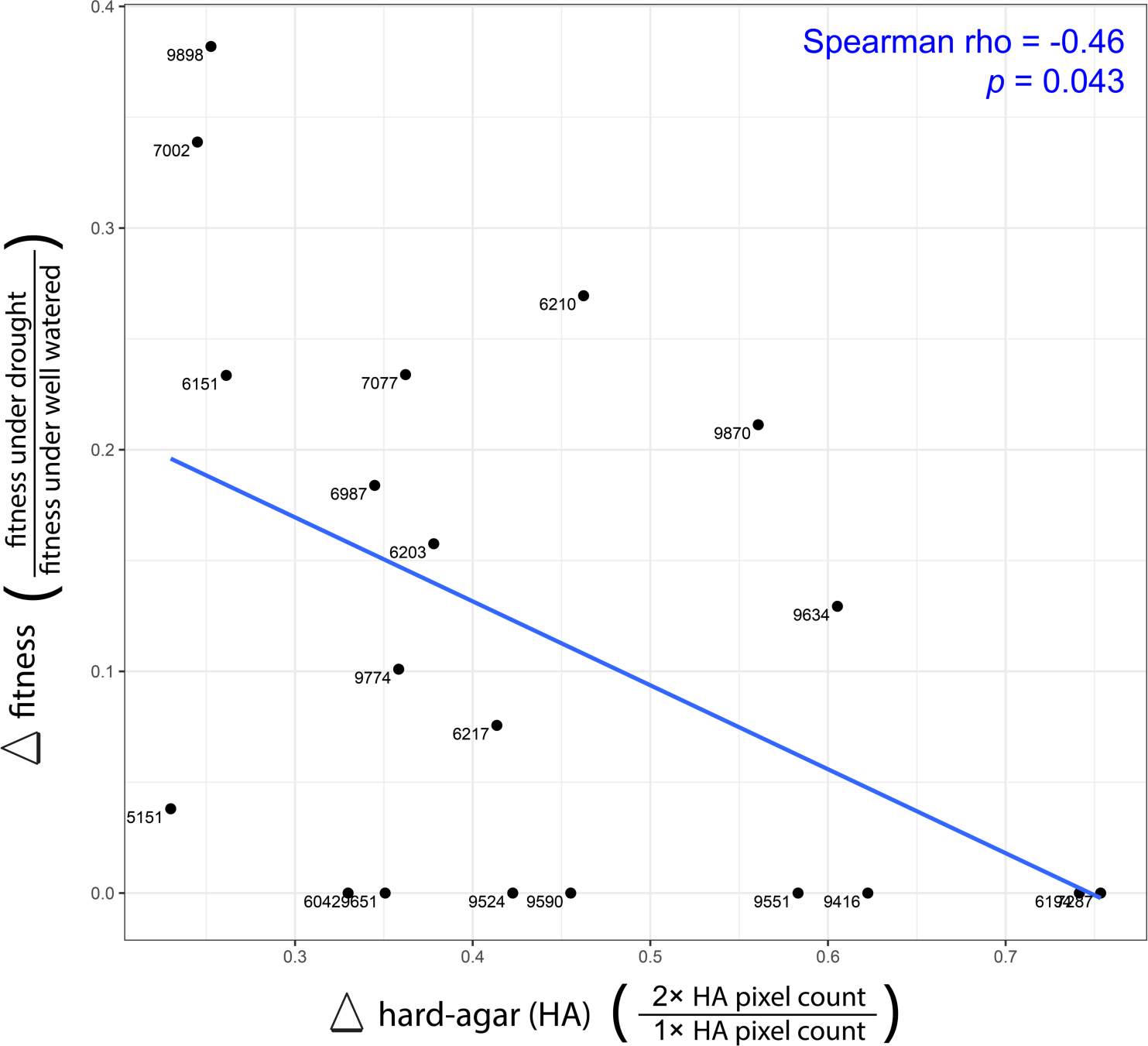
Associating hard agar’s (HA) impact on shoot size with plant fitness. Comparing the impact HA treatment has on shoot size of 20 different *Arabidopsis* accessions to the change in their fitness found under drought conditions in the field, as reported in (*27*).

**Figure 2-figure supplement 7.**
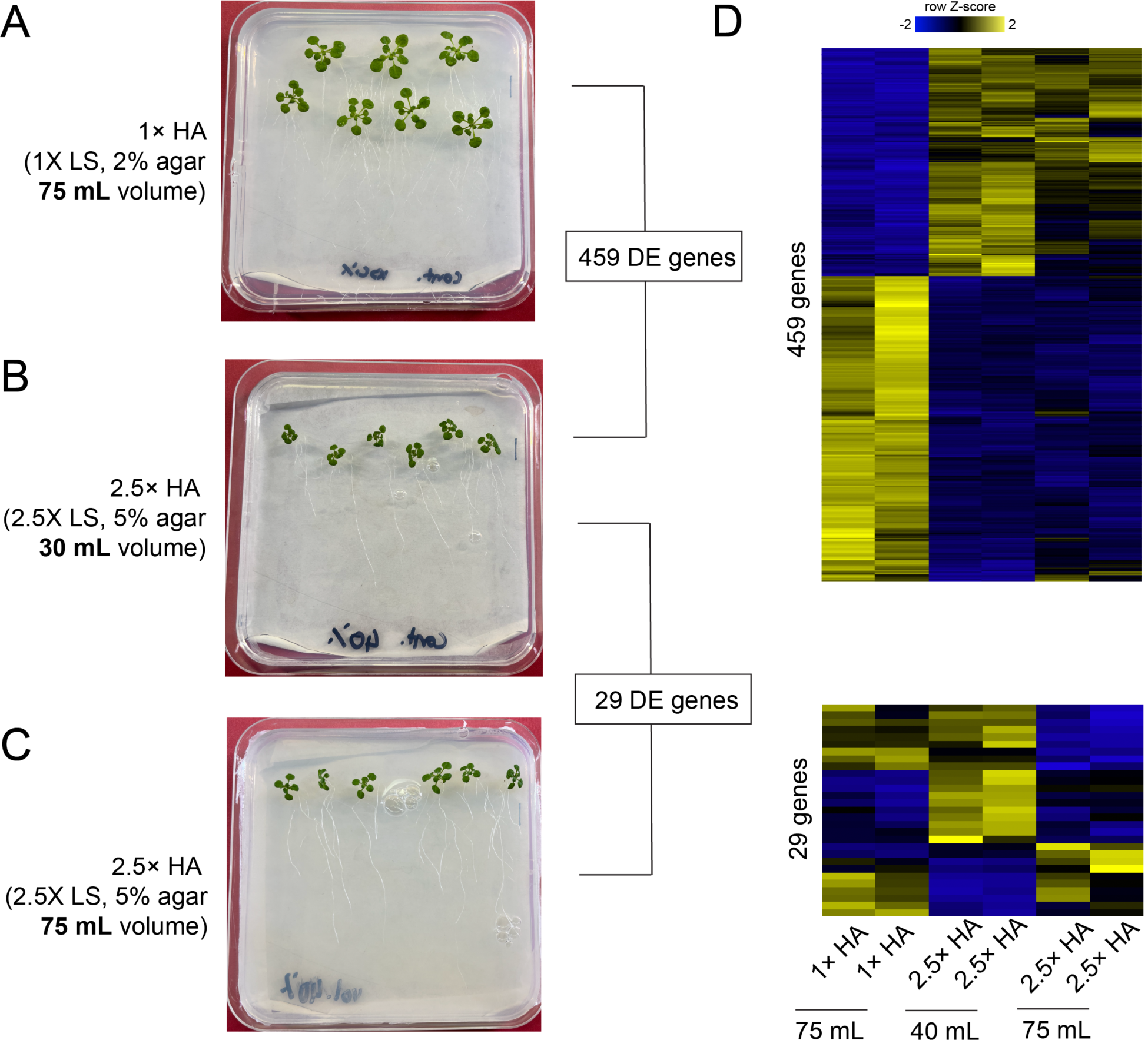
The volume of HA agar has minimal impact on gene expression. Image of *Arabidopsis* seedlings grown on either **(A)** 75 mL of 1ξ HA agar, **(B)** 30 mL of 2.5ξ HA agar, or **(C)** 75 mL of 2.5ξ HA agar. Number of genes found differentially expressed is reported for comparisons **(A)** and **(B),** as well as **(B)** and **(C)** (DESeq, adj. *p* < 0.01). Of the 29 genes found differentially expressed between **(B)** and **(C)**, 13 are found in comparison **(A)** and **(B)**. **(E)** Heatmap of genes found differentially expressed in either comparison.

